# Mian: Interactive Web-Based 16S rRNA Operational Taxonomic Unit Table Data Visualization and Discovery Platform

**DOI:** 10.1101/416073

**Authors:** Boyang Tom Jin

## Abstract

In recent years, there has been strong interest in examining the microbiome and its impact on human health and the environment. By leveraging modern sequencing technologies, investigators can quickly determine the composition of a given microbial sample. At the same time, the same investigations often yield an array of categorical and numerical metadata derived from the sequenced samples such as immunohistochemical measures or locality information. Understanding how the microbiome data is associated with this external metadata is essential in developing targeted treatments for chronic diseases or proposing bacteria-modulated host responses. While many R or Python libraries and command-line tools have been developed for specific analysis purposes, there are still relatively few tools to facilitate open-ended data exploration and hypothesis generation. Here we introduce Mian, an open-source web framework to interactively visualize or run a suite of statistical and feature selection tools on the microbiome to identify important taxonomic groups in the context of any provided categorical or numerical metadata. Visualizations include boxplots, correlation networks, and PCA or NMDS scatterplots. Tools include Fisher’s Exact Test, Boruta feature selection, alpha and beta diversity, and differential and correlational analysis. Mian supports multiple standard representations of the OTU table as input and optionally subsamples the data during the upload process. Users can also filter and aggregate the OTU table at different taxonomic levels and dynamically adjust analysis parameters to see how the visualizations, results, and statistical measures change in real-time. Mian is freely available at: miandata.org

## INTRODUCTION

The bacteria and archaea that form the microbiome have been the focus of many studies in recent years due to the influence they have on health and the environment (1, 2). In particular, research into the human microbiome has shown links between the presence of certain organisms and the development of certain human diseases such as inflammatory bowel disease and chronic obstructive pulmonary lung disease (COPD) (1). Studies done on the ocean microbiome have similarly uncovered the importance of marine microbes on key biogeochemical processes such as carbon and nutrient cycling (2, 3). Yet even today there is still much that remains undiscovered in the microbial world. By some estimates, less than 1% of all microbial species globally have been cultivated and categorized (4, 5). Because of this, it is thought that future profiling of microbial communities and better understanding of complex microbial relationships will fuel a diverse range of applications, from the development of new dietary interventions for chronic diseases (6), to introducing methods for more sustainable crop yields (7).

Traditionally, studies looking at the microbiome have primarily targeted the 16S rRNA owing to the universality of its distribution among bacteria and archaea, the relative low sequencing costs, and the availability of reference taxonomy databases such as Greengenes or SILVA (8, 9). A common approach to processing 16S rRNA sequencing data has been to group related sequences into operational taxonomic units (OTUs) which are then assigned to a reference taxonomy to determine the abundance and composition of a microbial community (5). Several software pipelines have been created to take the raw sequencing data and apply this OTU clustering analysis including Qiime and mothur (10, 11). As output, both of these pipelines can produce an OTU table, which is a two-dimensional matrix showing the abundance count of each OTU for each sample sequenced, along with the annotated OTU taxonomies. In order to provide a standardized method of storing these data along with sample and observational metadata, the BIOM file format was proposed to allow interoperability between different software platforms (12).

While data visualization is a crucial conduit in allowing users to detect patterns, trends, and relationships and to understand their underlying datasets, a core challenge in microbial analysis is to relate the changes the microbiome to contextual information regarding the samples themselves. For instance, when microbiome samples are sequenced from a human patient, researchers often also have access to a multitude of histological and pathophysiological data about the patient. Therefore, meaningful interpretation of the microbiome may only make sense when samples are grouped together according to this external metadata (13).

Another method of analyzing microbiome data is the use of data mining feature selection techniques, which can be used to identify statistically relevant organisms or taxonomic groups. These techniques are orthogonal to data visualizations as they rely on algorithms and statistical methods to facilitate hypothesis testing, as opposed to the data exploration capabilities that visualizations bring. It has been proposed that combining both data mining algorithms and visualizations will yield the most effective data discovery experience (14).

The most common method in applying data mining algorithms or generating visualizations is to use a common scripting language such as R or Python. However, this means that microbial researchers or clinicians who are not familiar with the computational aspects of microbial analysis are excluded from hypothesis testing and exploring data sets that they may have generated. Instead, they become reliant on bioinformaticians or statisticians to perform this type of analysis, increasing the overhead and turnaround time needed for studying the data. Moreover, although R or Python scripts can be automated or incorporated into broader bioinformatics pipelines, they are ultimately not as intuitive as a graphical user interface.

Here we introduce Mian, an interactive web-based data discovery platform that empowers users to examine the microbial community in the context of categorical and numerical metadata. Mian operates on OTU tables generated from existing OTU-picking software pipelines such as mothur and associated taxonomy annotations and user-provided sample metadata. Some of the key features of Mian are highlighted below:

- Generates interactive in-browser visualizations such as boxplots, scatterplots, and bar charts on microbial abundance, diversity measures, and provided sample metadata
- Supports principal component analysis (PCA) and non-metric multidimensional scaling (NMDS) to determine relatedness between distinct populations
- Runs powerful feature-selection algorithms and statistical tests such as Boruta/Random Forest and Fisher’s Exact Test to identify OTUs important in distinguishing between sample groupings
- Visualizations or feature-selection results respond in real-time with adjustments to the user-controllable input parameters
- OTUs can be grouped or filtered at different taxonomic levels during data analysis to show broad microbial patterns
- Provides statistical significance testing on select visualizations by providing p-values and FDR-corrected q-values
- Accepts standard OTU input formats including the BIOM file format and the mothur shared file

Mian is freely available for use at miandata.org. Mian is an open-source platform licensed under the MIT license with source code available at github.com/tbj128/mian.

## PROGRAM DESCRIPTION

### Pre-Requisites

Prior to using Mian, users are expected to have completed processing of their 16S rRNA sequencing data and have gone through an OTU-generation protocol such as the mothur MiSeq or 454 SOP (15, 16).

### Project Creation and Data Upload

Users first interact with Mian by creating an account. This account will store all of the project data that the user will upload and generate, allowing ease of use across multiple user sessions. Each project consists of an OTU table, the RDP, GreenGenes, or SILVA taxonomy annotations, and any metadata associated with the samples. All three pieces of information can either be uploaded in one single BIOM file or as separate tab-separated files. The BIOM file-format is recommended as it is a compact and widely used method to store and transfer OTU data between different data analysis software (12).

### Data Normalization

To avoid extraneous effects due to differences in sampling depth and to facilitate comparative analysis between different samples and groups, Mian visualizations and feature selection algorithms are designed to work against a normalized OTU table. There are three main normalization methods used in microbial ecology: rarefying, scaling, and transforming (17). Mian uses the rarefying method, which involves subsampling all samples to the same depth. While rarefying does reduce a certain amount of statistical power due to the removal of OTUs (17), rarefying reduces noise from OTUs which would not have been statistically relevant. This is a standard protocol used in many previous microbial studies and has been shown to be at least comparable to other normalization methods, although this remains a contested subject (18). Further OTU filtering based on user-configurable minimum count and prevalence thresholds is used in the feature selection tools. This is done to improve performance and increase the statistical power by excluding OTUs which will not likely be useful for further analysis.

Within Mian, users can either upload OTU data that is already subsampled or request Mian to subsample the data when creating a new project. By default, Mian will randomly subsample each sample to the overall lowest depth across all of the sample. Alternatively, users can also choose to define custom subsampling depths and filter out any samples falling below this depth. This latter approach is useful when the data contains outlier samples which would otherwise mislead downstream analysis.

The subsampled OTU table is used for all visualizations and feature selection tools with the exception of the rarefaction curve visualization which uses the original user-uploaded OTU table. The rarefaction curve allows a user to visually observe the number of OTUs selected as a function of subsampling depth. The user can then choose to adjust the subsampling depth if they observe the species diversity is too under-represented at the current subsampling depth.

### User Configurable Parameters

Each visualization or feature selection tool are shown as individual pages accessible from a dropdown menu. Within each page, users are presented with a series of filtering, aggregation, and analysis-specific parameter options that they can modify to suit their data exploration needs.

With the filtering option, users can choose to include or exclude OTUs and/or samples based on their associated taxonomic annotations and mappings. For instance, users can choose to only include OTUs from a single phylum in their analysis. Alternatively, users can decide to exclude any samples that have a particular metadata value. These options provide flexibility for the investigator to examine a subset of the data without having to reprocess the entire dataset. Feature selection modules have an additional layer of low-expression filtering. Only OTUs which meet a minimum user-configurable count and prevalence threshold will be considered as part of the analysis. By default, only OTUs whose abundance is greater than or equal to two in 10% or more samples will be kept for analysis. This is done in order to reduce the number of redundant computations on OTUs which will not produce any meaningful results.

With the aggregation option (“Taxonomic Level”), users can instruct Mian to collapse the OTU table at a higher taxonomic level by grouping and summing OTUs together according to the taxonomic annotations prior to applying the analysis. Aggregations can help establish trends or show patterns that occur across an entire family or phylum - information that otherwise may not have been available at just the OTU level. Aggregations can be particularly useful if the overall OTU table is extremely sparse by increasing statistical resolution to make more definitive conclusions.

Each individual tool will also have its own set of parameters which can be adjusted to suit each use case. Every time that a parameter is changed, Mian will immediately update the visualization or feature selection results to reflect the new state. This implementation allows users to more easily explore their dataset by quickly seeing how their analysis results change under different conditions and by removing any visual distractions such as page navigations.

## TOOLS SUMMARY

### Ordination Tool

Mian supports both principal component analysis (PCA) and non-metric multidimensional scaling (NMDS) for dimensionality reduction of the OTU table. The captured variation is presented on a two-dimension ordination plot. In the case of PCA, the user can choose to see which two principal components to view after viewing the overall variation represented by each of the first five principal components. In order to help highlight patterns and clustering within the dataset, users can also color the samples in the PCA plots based on a selected metadata attribute.

### Composition Tool

The composition of the OTU table can be represented at different taxonomic level groupings with bar and donut charts. These charts visually show the percentage abundance of each taxonomic group based on the mean of aggregate OTU counts across all samples. Users can also group the samples based on categorical metadata, which will generate a composition chart for each metadata value. This can, for example, allow the user to visually see the compositional differences between samples under different environmental conditions. These charts are also interactive, which encourages users to hover over different components to see details on the specific abundances values and taxonomic groups.

### Diversity Tool

The microbial community diversity analysis can be performed by calculating either the alpha diversity (measurement of the species diversity within a community) or beta diversity (measurement of the degree of differentiation of a community in relation to other communities). Both diversity measurements are presented as boxplots, which can be broken down based on a categorical metadata attribute for comparative analysis. If a breakdown is applied, Mian will also apply a Welch’s t-test to calculate whether the diversity differences are statistically significant. Welch’s t-test was selected as it is more robust than the Student’s t-test, allowing for unequal variances and sample sizes which can occur when metadata values are unevenly distributed across the samples. Each data point on the boxplots is also interactive, allowing users to hover over them and view details regarding the exact sample ID and calculated diversity value.

For alpha diversity, Mian allows users to choose between the three available diversity indices (Shannon, Simpson, and Inverse Simpson) that the underlying R ‘vegan’ library provides. In addition to the alpha diversity measure, species richness and Pielou’s measure of species evenness can also be rendered on the boxplots. For beta diversity, Mian also supports three types of diversity index measures (Bray-Curtis, Sorenson, and Whittaker). The Bray-Curtis index, based on the abundance counts, is recommended to show the dissimilarities between the different communities.

### Comparative Tools

#### Boxplots

Boxplots are the fundamental method of conveying numerical information within Mian. In addition to displaying diversity measurements, users can also render boxplots to show the abundance of a particular OTU or taxonomic group and also the mean or maximum OTU abundance per sample. This tool can help investigators visualize the differences between distinct sample groupings. For instance, in a study looking at the microbial samples collected from different lakes, investigators could look up whether there was an enrichment in the OTUs from the Firmicutes phylum between the different lakes. Alternatively, any numerical sample metadata can also be displayed on the boxplots. The Welch’s t-test is used to indicate whether the boxplot results are statistically significant.

#### Tree View

While boxplots are useful in exploring numerical attributes in relation to different communities, there is sometimes a need to compare sample groupings at a finer resolution. Within Mian, this can be accomplished by using the tree-view tool. This view generates a taxonomic tree based on the taxonomic assignments of the OTUs, up until a user configurable taxonomic level. The leaves of the tree show the average, maximum, or non-zero counts of each taxonomic group in relation to the sample grouping. This can help the user visually find taxonomic groups which are highly enriched in one sample grouping, which can then be targets of further investigation. Because of data display limitations, it is recommended to only perform this analysis at the class level or higher or to first filter for specific taxonomic groups.

#### Correlation Analysis

In microbial studies, correlations are popular and straightforward ways to determine relationships between two numerical variables. For example, one may want to look at correlations between microbial diversity versus gene expression data. Mian provides three ways to explore the correlational aspect: simple correlation visualization, correlation network analysis, and correlation selection.

In the most basic use case, users can choose to render a two-dimensional correlation scatterplot after specifying the two numerical metadata and/or OTU abundance measures they would like to be included in the correlation. Each sample on the correlation graph can also be colored or sized differently to allow additional categorical or numerical metadata to be included within the same analysis. A Pearson correlation coefficient and corresponding p-value is generated for user reference.

To quickly gain insights into the relationships between OTUs or taxonomic groups, Mian provides a weighted co-correlation network tool. This analysis involves computing the correlations between every pair of taxonomic elements and visualizing the results as a network. The width of each line in the network denotes the strength of the corresponding correlation. While this type of analysis is popular within the greater biological community, having been applied to cancer or evolutionary genetic analysis, it has also been successfully applied in the context of the microbial research although to a lesser extent. For example, previous studies have used co-occurrence networks to study the microbial dynamics in the soil environments (19). Co-correlation analysis is useful in determining the relationships within a complex community, such as identifying groups of organisms that may have commensal interactions with each other or, conversely, organisms which may be antagonistic to each other in a given environmental setting. Moreover, centralized modules within a correlation network may also hint at “keystone species”, organisms whose presence may be key to the stability of a community. Duran-Pinedo et al., for example, applied correlation network analysis to the oral biofilm to identify key bacterial modules within the oral community which could distinguish between healthy and disease subjects (20).

Alternatively, the correlation feature selection tool can be used to automatically identify OTUs or taxonomic group that correlate with a selected numerical metadata attribute. This is done by calculating the Pearson correlation coefficient, p-value, and FDR-corrected q-value for each OTU or taxonomic group and subsequently displaying the most statistically correlated OTUs or taxonomic groups to the user. This type of analysis can thus be used to determine the types of changes that can occur in a microbiome during disease progression or a host immune response. For example, by correlating microbial abundance against expression of genes from identified canonical pathways, Jangi et al. showed a positive association between the abundance of *Methanobrevibacter* and *Akkermansia* species and the genes implicated in the pathogenesis of multiple sclerosis (21). This type of correlational analysis can therefore be used a starting point to quickly assess whether there are any taxonomic shifts that warrant further investigation.

#### Fisher’s Exact Test

The Fisher’s exact test is used to test the statistical significance within a contingency table. In Mian, this tool examines the presence-absence of OTUs or taxonomy groups in the context of two pairwise categorical metadata values. This test would then select those OTUs which are shown to be statistically significant in discriminating between the two sample groupings. For example, consider a study where the samples were either “Control” or “Disease”. A contingency table could be constructed for each OTU such that the columns represent the presence and absence counts of the “Control” samples and the rows represent the presence and absence counts of the “Disease” samples. OTUs that have presence-absence proportions which are significantly different between the “Control” and “Disease” groups (according to their p-value and FDR-corrected q-value) would be highlighted in a tabular format for further investigation. This test can be especially useful when the OTU table is particularly sparse, since the test is able to utilize the full dataset and not just the non-zero values. Moreover, this test is also most applicable when the sample sizes are relatively small (*n* < 1000), although it is still correct for studies with larger samples (22). In either case, it is recommended to group the OTUs at a higher taxonomic level prior to running this test in order to increase the overall presence-absence counts, reduce computational cycles, and increase statistical resolution.

#### Random Forest/Boruta

Random forest classification algorithms are popular in the life science fields in part due to their ability to assign the importance to the variables that make up the classification. Random forests work particularly well with life science datasets since these datasets often contain more variables than samples and random forests can effectively handle this by training many decision trees across random subsets of the data (23). When applied to an OTU table, random forest algorithms can select for OTUs which are important in distinguishing between distinct sample groupings. For example, Sze et al. used Boruta feature selection with random forest analysis to identify 10 core OTUs, including known pathogens such as *Haemophilus influenzae*, that could discriminate between the samples taken from GOLD stage 4 COPD patients and those from healthy control individuals (24).

Within Mian, users can apply the random forest analysis to their OTU dataset and the resulting variable importance measure is used as input to the Boruta feature selection algorithm. Alternatively, Mian can also show the OTUs ranked by the random forest feature importance values as produced by the Python scikit-learn library. Boruta is an “all-relevant” feature selection method, which is ideal for selecting all OTUs that are relevant in discriminating between populations, as opposed to simply finding the non-redundant ones (25). By selecting for all relevant OTUs, one can start to explore patterns and commonalities between the selected OTUs such as shared metabolic processes or common environmental growth conditions.

#### Elastic Net

Mian supports a limited logistic and multinomial regression analysis for categorical metadata attributes using the R package GLMNet (26). GLMNet applies the elastic net regularization that linearly combines both lasso and ridge regularization penalties. Elastic net is particularly useful when the number of predictors outweigh the number of samples, such as the case of an OTU table. Elastic net also exposes groupings within the data, as it is able to select all members within a correlated group (27). On the UI, users can tune some basic hyperparameters to the GLMNet package in order to determine the OTUs or taxonomic groups that are most associated with certain outcomes such as a disease state or a particular phenotype. These OTUs can then be investigated in further downstream analysis. Depending on the use case, the user can choose to adjust the alpha parameter (elastic net mixing parameter) to vary the balance between the L1 or L2 regularization. On the UI, Mian abstracts away the majority of the tunable GLMNet parameters by using the default values or using fixed strategies, such as using cross-validation to determine the ideal lambda parameter. While this simplifies the flow for the user and allows users to quickly experiment with this model, it is recommended that users complete the analysis by replicating the results in R if further study is required or if additional visualizations are needed, such as the cross-validation curve.

#### Differential Selection

Using differential selection, Mian users can identify individual OTUs or taxonomic groups whose abundances are statistically different between two groups of samples. This is done by applying the Welch’s t-test to each candidate OTU (after filtering out low-expressed OTUs which would not result in meaningful results). This type of analysis is useful in understanding whether there are organisms which can differentiate between two disease states or two environmental habitats. For example, Fujimura et al. performed this type of between-groups comparison to determine what gastrointestinal microbial members afforded the best protection against allergens. Within their study, they found that *Lactobacillus* enrichment was associated with the adaptive immune response to respiratory insults (28). This type of selection is the categorical counterpart to the correlations selection, which instead selects OTUs based on comparisons with numerical metadata factors.

## PROGRAM IMPLEMENTATION

Mian consists of a web-based user-interface (written in JavaScript, HTML, and CSS) and a server-side data processing framework (written in Python and R). After the end-user defines their filtering, aggregating, and visualization parameters, the user-interface issues Ajax requests to the server to transform the underlying matrix data depending on the selected visualization or feature selection tool. By using the Python-based Flask microframework, Mian can leverage popular open-source statistics and machine learning libraries including “scikit-learn”, “scikit-bio”, and “numpy”. The server also interfaces with R libraries such as “vegan”, “Boruta”, and “glmnet” using the “RPy2” Python package. Visualizations are rendered on the user-interface using D3, an open-source JavaScript library.

Mian has been tested for compatibility with major browsers including Google Chrome, Safari, and Firefox. Testing was performed on a Macbook with 16GB of RAM running a 3.1 GHz Intel Core i7 processor and on an AWS EC2 Ubuntu m5.large instance with 8GB of RAM running 2.5GHz Intel Xeon Platinum processor.

## USE CASE EXAMPLE

In order to demonstrate Mian usage in real-world settings, we used raw data deposited in the DataDryad online repository from a previously published study by Sze et al. that examined the lung microbiome in individuals with COPD (24, 29). We processed the raw data in accordance with the mothur 454 SOP that was used by the original paper (16). For our case, we subsampled the dataset to 676 reads, and used version 132 of the SILVA reference to perform the alignment and version 16 of the RDP reference to assign the taxonomy, both of which are more recent than the reference alignments used by the original paper. The output files of the 454 SOP were uploaded directly to Mian along with sample metadata that labeled the samples as either “Control” or “COPD Gold 4”.

By using the alpha diversity tool, the visualization showed that the Shannon diversity index was significantly higher in the COPD samples than the control samples (P < 0.01). This observation is in-line with the study findings which showed that a decline in microbial diversity was associated with the emphysematous destruction that is hallmark of COPD disease progression.

The Boruta feature selection also confirmed eight OTUs as being relevant in distinguishing between the control and COPD samples. These OTUs included those from the *Prevotella, Streptococcus*, and *Flavobacterium* genera. Similar OTUs were also identified by the GLMNet and differential selection tools. As shown in Figure 1, we can then easily use the boxplot module to visually depict the differences in abundance of a particular taxonomic group.

**Figure 1.**
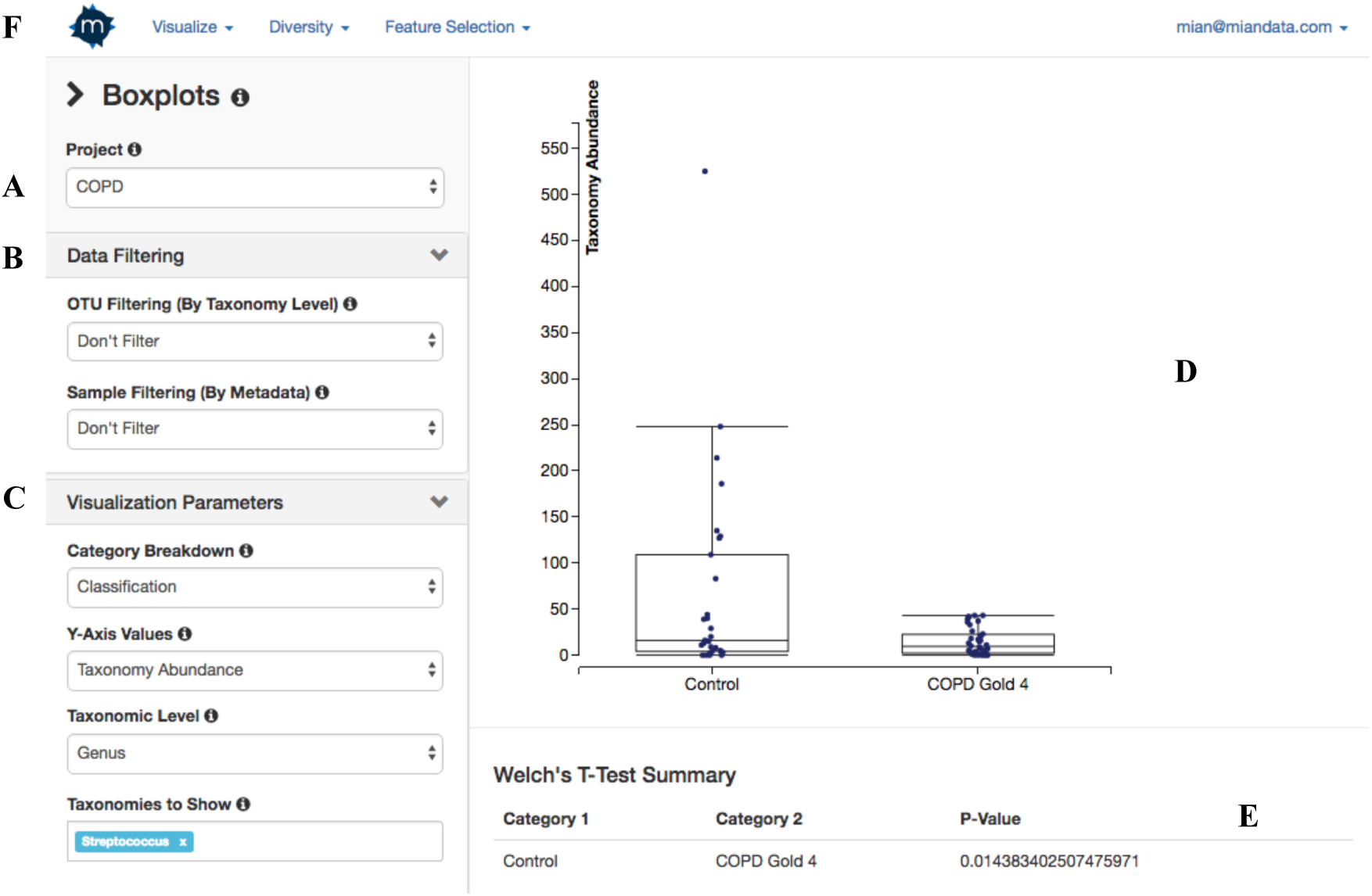
Boxplot tool layout in Mian; the layout is representative of most tools in Mian. **(A)** Project switcher changes the underlying source of the visualization or feature selection tool. **(B)** Data filtering parameters to selectively exclude or include OTUs/taxonomic groups and/or samples. In feature selection tools, there is an additional OTU filtering parameter based on minimum count and prevalence (not shown). **(C)** Visualization parameters that are specific to the tool which can change how samples are grouped and what values are displayed. Feature selection tools also allow adjustments to the input parameters of the underlying feature selection algorithms. **(D)** Main tool display area that shows either the desired visualization or a table of results. **(E)** Statistical significance of the visualization (if applicable). **(F)** Toolbar to access project details or other visualization or feature selection tools.

**Figure 2.**
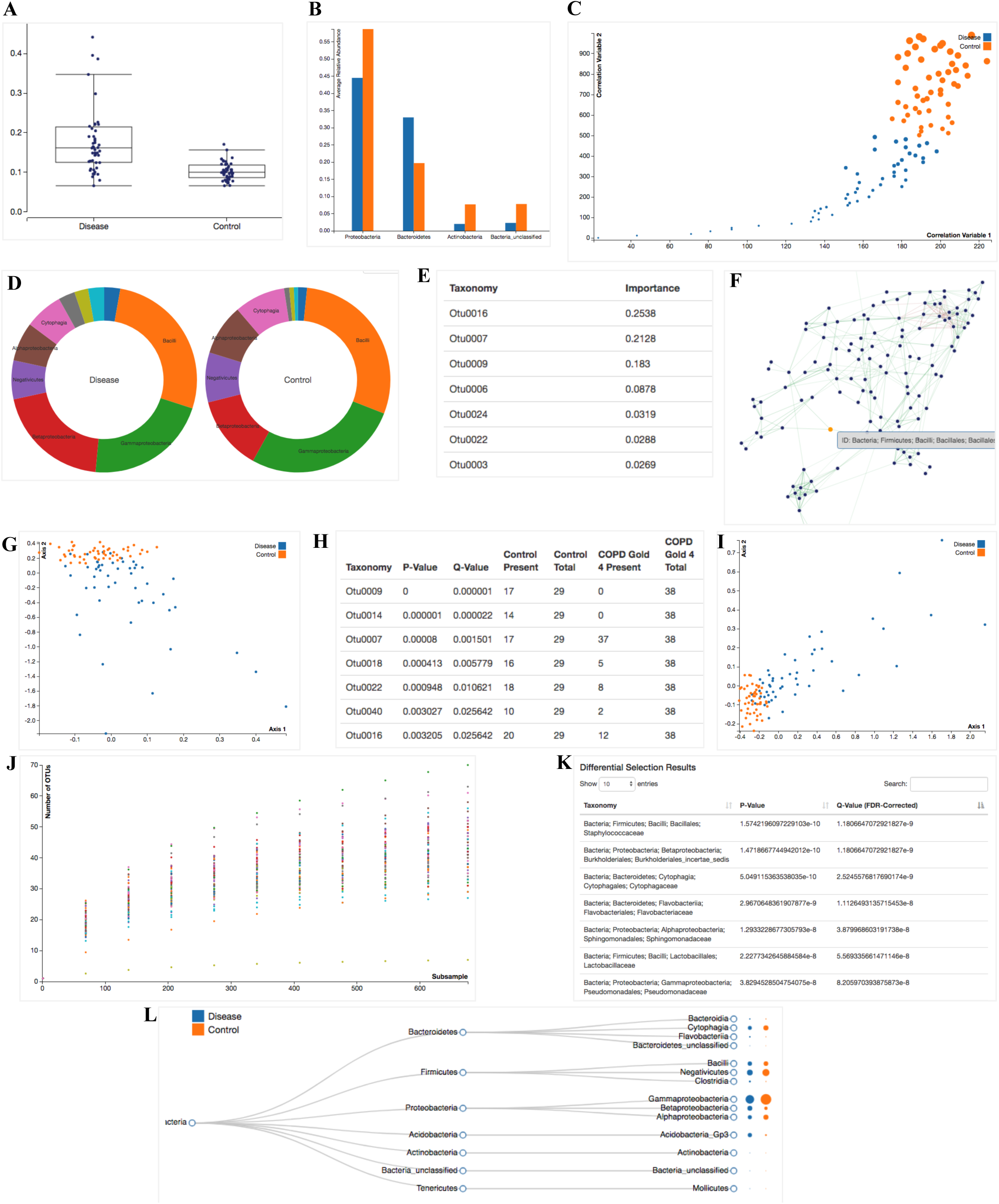
Examples of visualizations and feature selection results output available in Mian. **(A)** Beta diversity boxplot using the Bray-Curtis diversity index measure across two sample groups (not shown: similar boxplots are drawn for taxonomy abundance, numerical metadata, and alpha diversity). **(B)** Phylum-level composition comparison between two sample groups expressed as a bar chart. **(C)** Correlation scatterplot between taxonomy abundance and a numerical metadata attribute. **(D)** Class-level composition comparison between two sample groups expressed as a donut plot. **(E)** Random forest importance output at the OTU level. **(F)** Co-correlation network analysis grouped at the family level. **(G)** PCA scatterplot showing the first two principal components. **(H)** Fisher’s exact test to select for OTUs whose presence/absence can distinguish between control and COPD samples. **(I)** NMDS scatterplot colored according to their sample grouping. **(J)** Rarefaction curve showing OTU richness at different subsampling depths. **(K)** Differential selection results output after grouping the OTU table at the family level. **(L)** Tree view showing the relative abundance of each class with respect to each sample grouping.

When aggregated at the phylum level, the differential selection module showed that Proteobacteria phylum was an important differentiating factor between control and COPD samples (FDR < 0.002). Similarly, Sze et al. also found Proteobacteria to be the most significant driver of the difference between the control and the COPD samples. Both the PCA and NMDS plots also show a distinct separation between the microbiome communities of the control and COPD samples. Overall, the results obtained from Mian are comparable to the results from the COPD study. Some differences naturally arose due to differences in the subsampling depths and taxonomy reference files used.

## LIMITATIONS AND FUTURE DIRECTION

Because Mian is intended to be an interactive, real-time web-based platform, performance can become a bottleneck. The core data uploaded to Mian is a two-dimensional matrix representing the OTU table. As thousands of OTUs are typically generated through the upstream OTU selection process, this table rapidly grows in size with the number of samples which in turn consumes more computational and memory resources. Mian performs best when the OTU table is < 5MB (post-subsampling) - computationally expensive analysis may perform poorly on OTU tables larger than this size. For this reason, additional data filters based on minimum OTU count and prevalence on feature selection tools were implemented in order to reduce unnecessary computations on OTUs or taxonomic groups which would be poor targets of any further investigations due to their low expression.

Future iterations of Mian will look at using Map-Reduce libraries such as Apache Spark running on distributed cloud infrastructure in order to process large datasets in a more efficient manner. Mian will also look at introducing ways to more intuitively explore and visualize gene expression data, among other systems biology datasets, in the context of the microbiome community structure. In addition, we will look at augmenting the existing statistical and visualization tools with any tools that are found to be relevant to the microbial research community but currently missing in Mian.

## CONCLUSION

The microbiome is an ever-growing popular field of study due to its implications in a multitude of fields including medicine and the environment. In particular, microbial sequencing represents an important avenue of investigation in current human health related research as shifts in an individual’s microbiome have been associated with idiopathic diseases such as psoriasis and inflammatory bowel disease (30). With the costs of sequencing also continuing to decline, microbial sequencing is becoming more and more popular and accessible which will inevitably lead to a concomitant increase in the amount of data produced. Nevertheless, microbial sequencing is still a relatively young field. Many studies are still exploratory in nature and try to infer general patterns based on differential analysis. Therefore, it is critical to have tools that can facilitate data exploration in order to drive hypothesis generation and gain insights into these large microbial datasets. As part of this explorative narrative, these tools should also juxtapose any numerical and categorical metadata available on the microbial samples in order to drive research questions and shape future experiments.

Mian fulfills this need by packaging several commonly used visualization, feature selection, and diversity tools into a user-friendly unified web interface. This design enables users without any programming or scripting background to upload their data sets and immediately generate boxplots or run a random forest feature selection algorithm without any data manipulation required. While tools have been created to generate visualizations based on microbial data sets such as EMPeror (13), Mian extends this concept further with novel visualizations such as the tree view to explore the relationship between external metadata and the microbial community. Furthermore, Mian emphasizes the use of machine learning and data mining feature selection algorithms to discover individual organisms or groups of organisms that may together contribute to a particular response in a population or result in a disease state. We believe the results derived from Mian will allow researchers to better integrate multiple datasets and more quickly explore different hypotheses to identify the ones that are most promising to pursue. Ultimately, Mian seeks to help researchers set the direction of their research and to remove technical hurdles in the data exploration process.

## REFERENCES

1. Shreiner AB, Kao JY, Young VB. The gut microbiome in health and in disease. Current opinion in gastroenterology. 2015 Jan;31(1):69.

2. Moran MA. The global ocean microbiome. Science. 2015 Dec 11;350(6266):aac8455.

3. White III RA, Callister SJ, Moore RJ, Baker ES, Jansson JK. The past, present and future of microbiome analyses. Nature Protocols. 2016 Nov;11(11):2049.

4. Locey KJ, Lennon JT. Scaling laws predict global microbial diversity. Proceedings of the National Academy of Sciences. 2016 May 24;113(21):5970–5.

5. Chen W, Zhang CK, Cheng Y, Zhang S, Zhao H. A comparison of methods for clustering 16S rRNA sequences into OTUs. PloS one. 2013 Aug 13;8(8):e70837.

6. Berg G, Grube M, Schloter M, Smalla K. Unraveling the plant microbiome: looking back and future perspectives. Frontiers in microbiology. 2014 Jun 4;5:148.

7. Bäckhed F, Fraser CM, Ringel Y, Sanders ME, Sartor RB, Sherman PM, Versalovic J, Young V, Finlay BB. Defining a healthy human gut microbiome: current concepts, future directions, and clinical applications. Cell host & microbe. 2012 Nov 15;12(5):611–22.

8. Větrovský T, Baldrian P. The variability of the 16S rRNA gene in bacterial genomes and its consequences for bacterial community analyses. PloS one. 2013 Feb 27;8(2):e57923.

9. Balvociute M, Huson DH. SILVA, RDP, Greengenes, NCBI and OTT—how do these taxonomies compare?. BMC genomics. 2017 Mar;18(2):114.

10. Caporaso JG, Kuczynski J, Stombaugh J, Bittinger K, Bushman FD, Costello EK, Fierer N, Pena AG, Goodrich JK, Gordon JI, Huttley GA. QIIME allows analysis of high-throughput community sequencing data. Nature methods. 2010 May;7(5):335.

11. Schloss PD, Westcott SL, Ryabin T, Hall JR, Hartmann M, Hollister EB, Lesniewski RA, Oakley BB, Parks DH, Robinson CJ, Sahl JW. Introducing mothur: open-source, platform-independent, community-supported software for describing and comparing microbial communities. Applied and environmental microbiology. 2009 Dec 1;75(23):7537–41.

12. McDonald D, Clemente JC, Kuczynski J, Rideout JR, Stombaugh J, Wendel D, Wilke A, Huse S, Hufnagle J, Meyer F, Knight R. The Biological Observation Matrix (BIOM) format or: how I learned to stop worrying and love the ome-ome. GigaScience. 2012 Dec;1(1):7.

13. Vázquez-Baeza Y, Pirrung M, Gonzalez A, Knight R. EMPeror: a tool for visualizing high-throughput microbial community data. GigaScience. 2013 Dec;2(1):16.

14. Shneiderman B. Inventing discovery tools: combining information visualization with data mining. Information visualization. 2002 Mar;1(1):5–12.

15. Kozich JJ, Westcott SL, Baxter NT, Highlander SK, Schloss PD. Development of a dual-index sequencing strategy and curation pipeline for analyzing amplicon sequence data on the MiSeq Illumina sequencing platform. Applied and environmental microbiology. 2013 Jun 21:AEM-01043.

16. Schloss PD, Gevers D, Westcott SL. (2011). Reducing the effects of PCR amplification and sequencing artifacts on 16S rRNA-based studies. PloS ONE. 6:e27310.

17. Weiss S, Xu ZZ, Peddada S, Amir A, Bittinger K, Gonzalez A, Lozupone C, Zaneveld JR, Vázquez-Baeza Y, Birmingham A, Hyde ER. Normalization and microbial differential abundance strategies depend upon data characteristics. Microbiome. 2017 Dec;5(1):27.

18. Weiss SJ, Xu Z, Amir A, Peddada S, Bittinger K, Gonzalez A, Lozupone C, Zaneveld JR, Vazquez-Baeza Y, Birmingham A, Knight R. Effects of library size variance, sparsity, and compositionality on the analysis of microbiome data. PeerJ PrePrints; 2015 Jun 6.

19. Barberán A, Bates ST, Casamayor EO, Fierer N. Using network analysis to explore co-occurrence patterns in soil microbial communities. The ISME journal. 2012 Feb;6(2):343.

20. Duran-Pinedo AE, Paster B, Teles R, Frias-Lopez J. Correlation network analysis applied to complex biofilm communities. PloS one. 2011 Dec 7;6(12):e28438.

21. Jangi S, Gandhi R, Cox LM, Li N, Von Glehn F, Yan R, Patel B, Mazzola MA, Liu S, Glanz BL, Cook S. Alterations of the human gut microbiome in multiple sclerosis. Nature communications. 2016 Jun 28;7:12015.

22. Connelly LM. Fisher’s exact test. MedSurg Nursing. 2016;25(1):58.

23. Touw WG, Bayjanov JR, Overmars L, Backus L, Boekhorst J, Wels M, van Hijum SA. Data mining in the Life Sciences with Random Forest: a walk in the park or lost in the jungle?. Briefings in bioinformatics. 2012 Jul 10;14(3):315–26.

24. Sze MA, Dimitriu PA, Suzuki M, McDonough JE, Campbell JD, Brothers JF, Erb-Downward JR, Huffnagle GB, Hayashi S, Elliott WM, Cooper J. Host response to the lung microbiome in chronic obstructive pulmonary disease. American journal of respiratory and critical care medicine. 2015 Aug 15;192(4):438–45.

25. Kursa MB, Rudnicki WR. Feature selection with the Boruta package. J Stat Softw. 2010 Sep 16;36(11):1–3.

26. Friedman JH, Hastie TJ, Tibshirani RJ. glmnet: lasso and elastic-net regularized generalized linear models, 2010b. URL http://CRANR-project.org/package=glmnet. R package version.:1-.

27. Zou H, Hastie T. Regularization and variable selection via the elastic net. Journal of the Royal Statistical Society: Series B (Statistical Methodology). 2005 Apr 1;67(2):301–20.

28. Fujimura KE, Demoor T, Rauch M, Faruqi AA, Jang S, Johnson CC, Boushey HA, Zoratti E, Ownby D, Lukacs NW, Lynch SV. House dust exposure mediates gut microbiome Lactobacillus enrichment and airway immune defense against allergens and virus infection. Proceedings of the National Academy of Sciences. 2014 Jan 14;111(2):805–10.

29. Sze MA, Dimitriu PA, Suzuki M, McDonough JE, Campbell JD, Brothers JF, Erb-Downward JR, Huffnagle GB, Hayashi S, Elliott WM, Cooper J, Sin DD, Lenburg ME, Spira A, Mohn WW, Hogg JC (2015) Data from: The host response to the lung microbiome in Chromic Obstructive Pulmonary Disease. Dryad Digital Repository. https://doi.org/10.5061/dryad.2p66n

30. Cho I, Blaser MJ. The human microbiome: at the interface of health and disease. Nature Reviews Genetics. 2012 Apr;13(4):260.

